# Insomnia subtypes have differentiating deviations in brain structural connectivity

**DOI:** 10.1101/2023.11.01.565094

**Authors:** T. Bresser, T.F. Blanken, S. C. de Lange, J. Leerssen, J.C. Foster-Dingley, O. Lakbila-Kamal, R. Wassing, J.R. Ramautar, D. Stoffers, M.P. van de Heuvel, E.J.W. van Someren

**Affiliations:** Netherlands Institute for Neuroscience, Department of Sleep and Cognition, Amsterdam, Netherlands; Vrije Universtiteit Amsterdam, Department of Integrative Neurophysiology, Amsterdam, Netherlands; Department of Complex Trait Genetics, Center for Neurogenomics and Cognitive Research, Amsterdam Neuroscience, Vrije Universiteit Amsterdam, Amsterdam, The Netherlands; University of Amsterdam, Department of Psychological Methods, Amsterdam, Netherland; Department of Psychiatry, Vrije Universtiteit Amsterdam, Amsterdam UMC, Amsterdam, The Netherlands; Sleep and Circadian Research, Woolcock Institute of Medical Research, Macquarie University Sydney, NSW, Australia; School of Psychological Sciences, Faculty of Medicine, Health and Human Sciences, Macquarie University, Sydney, NSW, Australia; N=You Neurodevelopmental Precision Center, Amsterdam Neuroscience, Amsterdam Reproduction and Development, Amsterdam UMC, Amsterdam, The Netherlands; Child and Adolescent Psychiatry and Psychosocial Care, Emma Children’s Hospital, Amsterdam UMC, Vrije Universiteit Amsterdam, Amsterdam, the Netherlands; Spinoza Centre for Neuroimaging, Amsterdam, The Netherlands; Department of Child and Adolescent Psychiatry and Psychology, Amsterdam UMC location Vrije Universiteit Amsterdam, Amsterdam, The Netherlands

## Abstract

**Objective:** Insomnia disorder is the most common sleep disorder. A better understanding of insomnia-related deviations in the brain could inspire better treatment. Insufficiently recognized heterogeneity within the insomnia population could obscure involved brain circuits. The present study investigated whether structural brain connectivity deviations differ between recently discovered and validated insomnia subtypes.

**Methods:** Structural and diffusion weighted 3-Tesla MRI data of four independent studies were harmonized. The sample consisted of 73 controls without sleep complaints and 204 participants with insomnia grouped into five subtypes based on their fingerprint of personality and mood traits assessed with the Insomnia Type Questionnaire. Linear regression correcting for age, sex, and brain volume evaluated group differences in structural connectivity strength, indicated by fractional anisotropy and mean diffusivity, and evaluated within two different atlases.

**Results:** Insomnia subtypes showed differentiating profiles of deviating structural connectivity which moreover concentrated in different functional networks. Permutation testing against randomly drawn heterogeneous subsamples indicated significant specificity of deviation profiles in four of the five subtypes: *highly distressed* (p=0.019)*, moderately distressed reward insensitive* (p=0.014)*, slightly distressed low reactive* (p=0.006) and *slightly distressed high reactive* (p=0.006).

**Conclusions:** Our results provide a first indication that different insomnia subtypes exhibit distinct profiles of deviations in structural brain connectivity. Subtyping of insomnia could be essential for a better understanding of brain mechanisms that contribute to insomnia vulnerability.

## Introduction

Insomnia disorder is a common sleep disorder affecting approximately 10% of the adult European population (1,2). Insomnia patients suffer from persistent difficulty falling asleep, staying asleep and/or early morning awakening with subjectively impairing day-time functioning (3). Insomnia disorder has severe consequences, including an increased risk of cardiovascular disorders (4), obesity (5) and mental disorders (6,7). Cognitive behavioral therapy for insomnia (CBT-I) can alleviate the burden of insomnia complaints (8,9) and contribute to prevention of other mental disorders (10–12). However, CBT-I does not bring sufficient relief for all patients, even if complemented with hypnotics (8,9). In order to innovate and improve treatment, we need a better understanding of brain circuits involved in insomnia vulnerability.

Clues to altered brain networks can be pursued by studying structural connectivity between brain regions at the macroscopic scale using neuroimaging techniques (13). Reviews of structural and functional imaging studies comparing people suffering from insomnia disorder and people without sleep complaints have suggested a potential involvement of the default mode network and salience network in insomnia (14–18). Nevertheless, explained variance and consistency across studies has been limited (19,20). It has been proposed that inconsistency in neuroimaging results could in part be due to unrecognized heterogeneity within the affected population: differently distributed deviations in brain structure or function might present as the same disease phenotype (21,22).

If more homogeneous subtypes could be recognized within such a heterogeneous population, corresponding deviations in structural connectivity may be determined with better consistency. How to define different insomnia subtypes is an ongoing discussion. Various insomnia subtype classifications have focused on dominant sleep complaints (3,23–26). Unfortunately, sleep feature-related classifications may not be that robust (23,25). More recently a bottom-up, data driven approach revealed five more robust subtypes of insomnia disorder. Instead of focusing on sleep features only, the subtypes could be distinguished based on their level of distress as well as their unique profile of personality and mood traits (22). These subtypes can be assessed with the Insomnia Type Questionnaire (ITQ) (22). It can be hypothesized that ITQ-based insomnia subtypes also differ with respect to deviations in the brain circuits involved in these distinguishing personality and mood traits.

The present study therefore aimed to compare structural connectivity of the five ITQ-based insomnia subtypes. Based on brain region-to-function mappings reported in the literature (27), and moreover based on subtype-specific personality and mood traits, we selected frontal, orbitofrontal and temporal brain regions for inclusion in analyses. White matter microstructure of structural connections between the selected regions was assessed using fractional anisotropy (FA) and mean diffusivity (MD) (28,29). Analysis of 204 subtyped people with insomnia disorder and 73 people without sleep complaints, provides a first indication that insomnia subtypes can have different structural connectivity deviation profiles and sometimes even opposing deviations. We further contextualized our findings by identifying the major functional networks that were most affected for each subtype.

## Methods

### Participants

Data were acquired during four studies performed between 2014 and 2021 by the Sleep & Cognition group at the Netherlands Institute for Neuroscience, Amsterdam (30–32). Participants were recruited through the Netherlands Sleep Registry (www.slaapregister.nl), advertisement, and media. Across all studies, applicants were eligible if they were between 18 and 70 years of age. Screening occurred via telephone, online questionnaires and subsequent intake interview. Depending on the original study sample, insomnia disorder was diagnosed in accordance to the International Classification of Sleep Disorders, third edition (24) and Diagnostic and Statistical Manual of Mental Disorders Fourth (33) or Fifth Edition (3). Exclusion criteria for applicants varied between original studies but consistently were a current diagnosis of severe sleep apnea, severe restless leg syndrome, narcolepsy, any other severe neurological, psychiatric, or somatic disorders, current shift work, or any MRI contraindication such as non-MR-compatible metal implants, claustrophobia, or pregnancy (see supplement for additional details). In addition, three studies recorded a polysomnogram which was inspected for undiagnosed severe sleep disorders other than insomnia. All studies were approved by the ethical board of the VU Medical Center or the University of Amsterdam. Written informed consent was obtained from all participants.

### Measures

Sample characteristics were described using the insomnia severity index (ISI, range 0 - 28)(34), Pittsburgh Sleep Quality Index (PSQI, range 1 - 21) (35), inventory of depressive symptomatology – self report (IDS-SR, range 0 - 84) (36), Hospital Anxiety and Depression Scale (HADS, anxiety and depression subscale both range 0 - 21) (37). In these questionnaires, higher scores indicate more severe symptoms. In participants with a diagnosis of insomnia disorder we determined the insomnia type using the ITQ (22). T1-weighted and diffusion-weighted images were acquired using two Philips Achieva 3T scanners (see supplement for MRI acquisition details).

### Structural connectome reconstruction

Preprocessing and reconstruction of structural connectivity networks was performed separately for each sample. T1w scans were preprocessed and segmented using ‘recon-all’ function of FreeSurfer (38) stable version 6.0.1. Connectivity Analysis TOolbox (39) (CATO, version 3.1.2) and FMRIB Software Library (40) (FSL version 6.0.4) were used to preprocess diffusion weighted data and reconstruct structural connectivity matrices. For network reconstruction we used the fine-grained Cammoun sub-parcellation of the Desikan-Killiany atlas consisting of 114 cortical regions (39,41). To validate our findings and include subcortical regions, we also used the Desikan-Killiany combined cortical and subcortical atlas as present in FreeSurfer (42) consisting of 82 regions. For each atlas, the weight of a connection between regions was computed as the weighted average FA or MD value over all voxels that the streamlines passed. To ensure sufficient data points for each connection, analyses included only connections that were present in at least 70% of the subjects (43). To account for multisite and multi-sample effects we harmonized the connection weights between all brain regions for sample differences using the ‘NeuroCombat’ R-package version 1.0.13 (https://github.com/Jfortin1/ComBatHarmonization) in R (44) version 4.0.4 (see supplement for more details).

### Regions-of-interest selection

The fact that insomnia subtypes differ with respect to their profile of personality and mood traits allowed for a data-driven selection of brain regions of interest representing these traits (figure 1A). To do so, we made use of the Neuro Knowledge Engine (NKE) (27), a functional neuroimaging database linking brain structures to function. NKE consists of functional terms and brain activation coordinates extracted from functional neuroimaging studies. By combining functional terms, NKE describes clusters that represent neurobiological domains e.g., arousal. We matched the mood and personality traits that distinguish ITQ-based insomnia subtypes with NKE-functional terms to identify NKE-neurobiological domains where insomnia subtypes differ (see supplement for a list of the used terms). NKE-neurobiological domains were included if at least 40% of the NKE-functional terms matched terms distinguishing insomnia subtypes. Next, the brain coordinate data of included NKE-neurobiological domains were registered to atlas space and the frequency of insomnia subtype-related NKE-neurobiological domains per atlas region was determined. NKE-functional terms can be clustered with arbitrary granularity into as little as 2, up to as much as 50, different NKE-neurobiological domains. We focused on the finer granularity of neurobiological domains and repeated the steps described above when NKE-functional terms were clustered in 25 up to 50 NKE-neurobiological domains. Iterating over the different NKE clustering granularities allowed us to build a heat map indicating the involvement of brain regions in insomnia (figure s1). We selected regions-of-interest that linked at least 10 times to any neurobiological domains associated with insomnia subtype traits.

**Figure 1.**
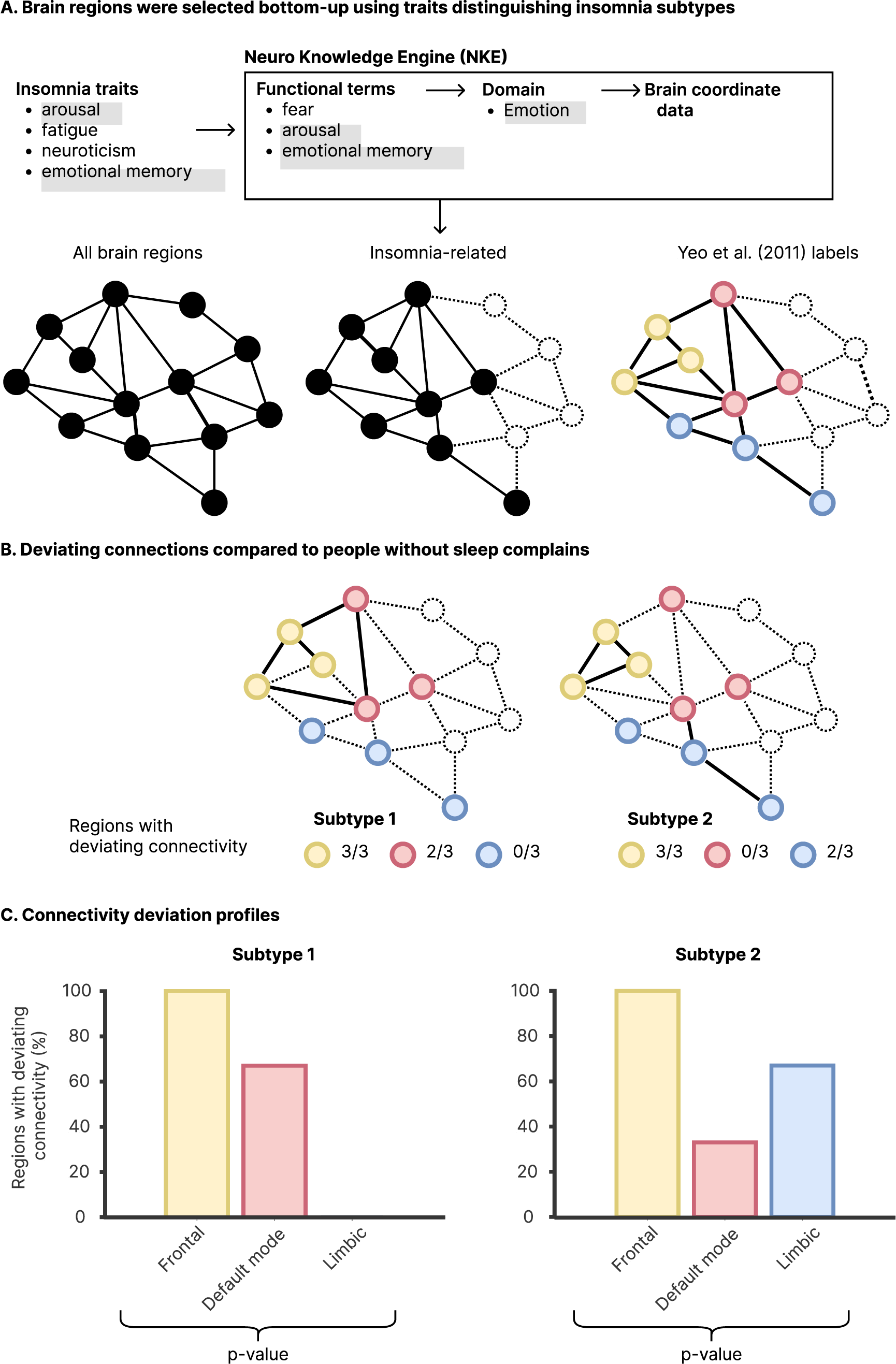
Procedure of brain region selection and obtaining connectivity deviation profiles. A) Brain regions involved in traits distinguishing insomnia subtypes were selected using the Neuro Knowledge Engine database and functionally annotated according to the Yeo *et al.* (2011) intrinsic functional resting-state networks. B) Examples of deviating (solid lines) and non-deviating (dashed lines) connections of insomnia subtypes 1 and 2 compared to people without sleep complaints. C) Example connectivity deviation profiles based on the proportion brain regions with deviating structural connectivity in each functional network. The p-value indicating significance of the specificity of the connectivity deviation profile of each subtype is obtained by permutation testing with drawing groups randomly from all heterogeneous insomniacs irrespective of their subtype, while keeping the control group the same.

### Deviating structural connectivity

After selecting brain regions involved in traits distinguishing insomnia subtypes (figure 1A), connection-wise deviations in white matter microstructure were assessed twice: based on FA and based on MD. For every connection, group differences in the standardized deviation of FA and MD were estimated by running two multivariate regression models on z-transformed data including the covariates age, sex and brain volume. A first model compared the heterogeneous insomnia group combining all subtypes, with the group of controls without sleep complaints. A second model compared all five insomnia subtypes as separate groups with people without sleep complaints. Subsequent analyses focused on ‘deviating connections’, defined as all deviations exceeding a primary (t-statistic) threshold of |t| >= 2 (figure 1B).

### Functional annotation and connectivity deviation profiles

The resulting matrices with deviating structural connections between the selected brain regions were functionally annotated to evaluate whether insomnia subtypes differed in the functional systems involved. Brain regions were annotated based on the seven intrinsic functional resting-state networks described by Yeo *et al.* (45). Next, we calculated the percentage of regions with deviating connectivity within each functional network by dividing the number of regions with at least one deviating connection exceeding a t-statistic |t|>=2, by the number of regions present in the functional network (figure 1B-C). This approach provided, both for the heterogeneous insomnia group and for each subtype, a profile of the proportion brain regions with deviating structural connectivity in each functional network.

### Connectivity deviation profile-based statistics

Permutation testing of the ‘connectivity deviation profiles’ was used to evaluate subtype specificity. This approach tests the probability of obtaining similar results if we would take random sub-samples of the total heterogeneous insomnia sample to compare with the group of controls without sleep complaints. We ran 10 000 permutations in which subtype labels were shuffled across participants with insomnia while taking covariates into account (46). In short (see also figure 1B-C), for each permutation we repeated the analysis steps described above by estimating connection-wise deviations in the randomly shuffled subtype groups and computing the number of brain regions with deviating structural connectivity in each annotated functional network. Based on the estimated null-distributions, p-values for the connectivity deviation profiles were obtained by calculating how often at least the same number of brain regions with deviating connectivity were present for all five functional network annotations in the randomly labeled groups. The p-values represent the likelihood of finding the subtype-specific connectivity deviation profile in randomly labeled subsamples of heterogeneous insomnia. Unless stated otherwise, all analyses were performed in MATLAB 2019b. Figures use an accessible qualitative color scheme (47). An adapted version of the matlab script ‘schemaball’ was used to obtain circular representations of deviating connections (https://github.com/GuntherStruyf/matlab-tools/blob/master/schemaball.m).

## Results

### Participants

The total sample consisted of a reference group of n=73 people without sleep complaints and n=204 people suffering from insomnia. Within the insomnia sample n=31 were subtyped as *highly distressed*, n=89 as *moderately distressed reward sensitive*, n=30 as *moderately distressed reward insensitive*, n=34 as *slightly distressed high reactive* and n=20 as *slightly distressed low reactive*, as described by Blanken et al (22). Table 1 shows all demographics.

**Table 1.**
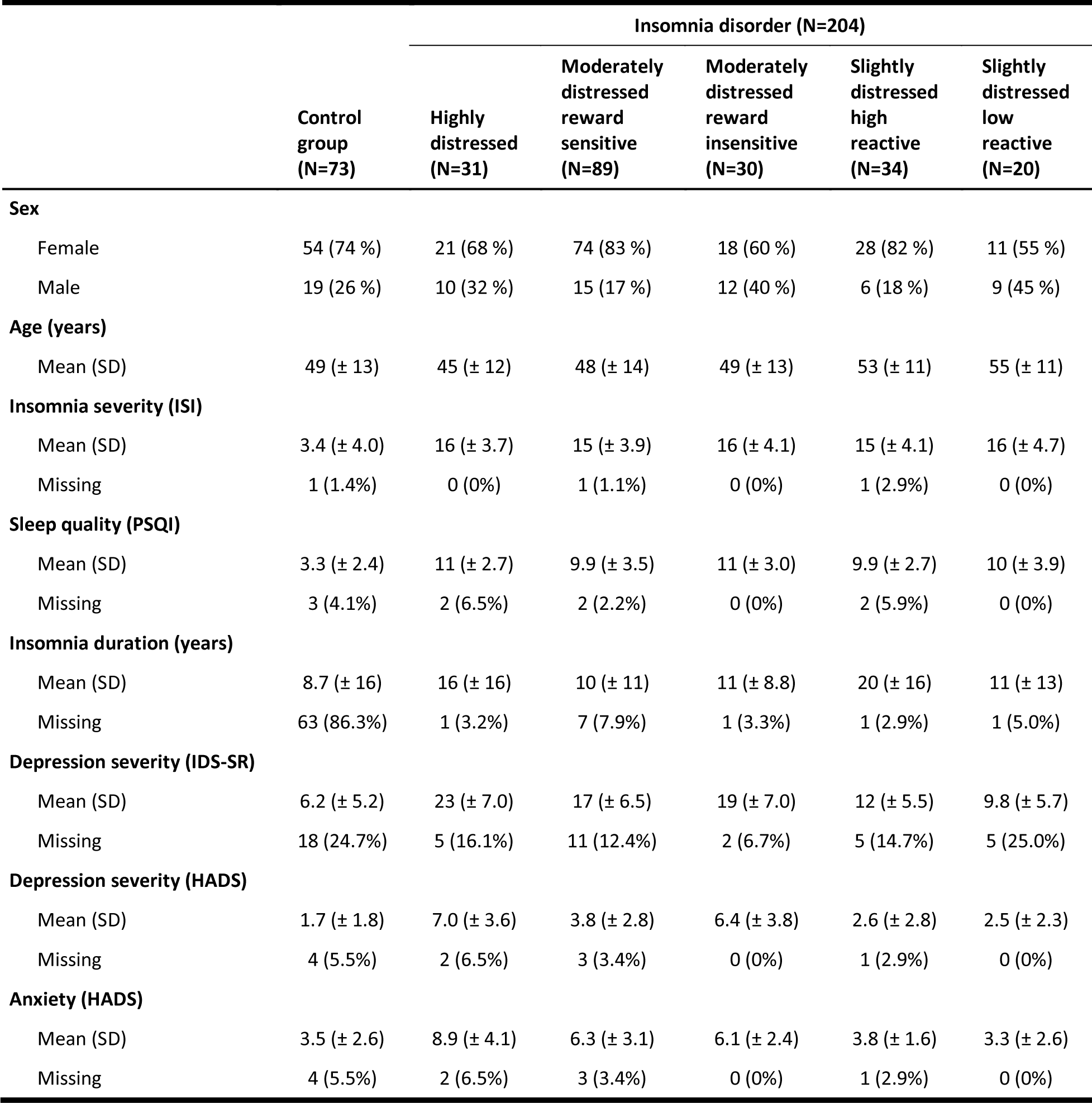
Sample characteristics.

### Insomnia subtype traits map to frontal, orbitofrontal and temporal brain regions

NKE was used to map mood and personality traits distinguishing insomnia subtypes to the Cammoun sub-parcellation of the Desikan-Killiany cortical atlas (41). The resulting heat map (figure s1) indicated involvement of frontal, orbitofrontal and temporal regions. Based on this heat map, 43 regions-of-interest (table s2) that linked at least 10 times to any NKE-neurobiological domain associated with insomnia subtype traits were included (Figure 2A) in subsequent analyses.

**Figure 2.**
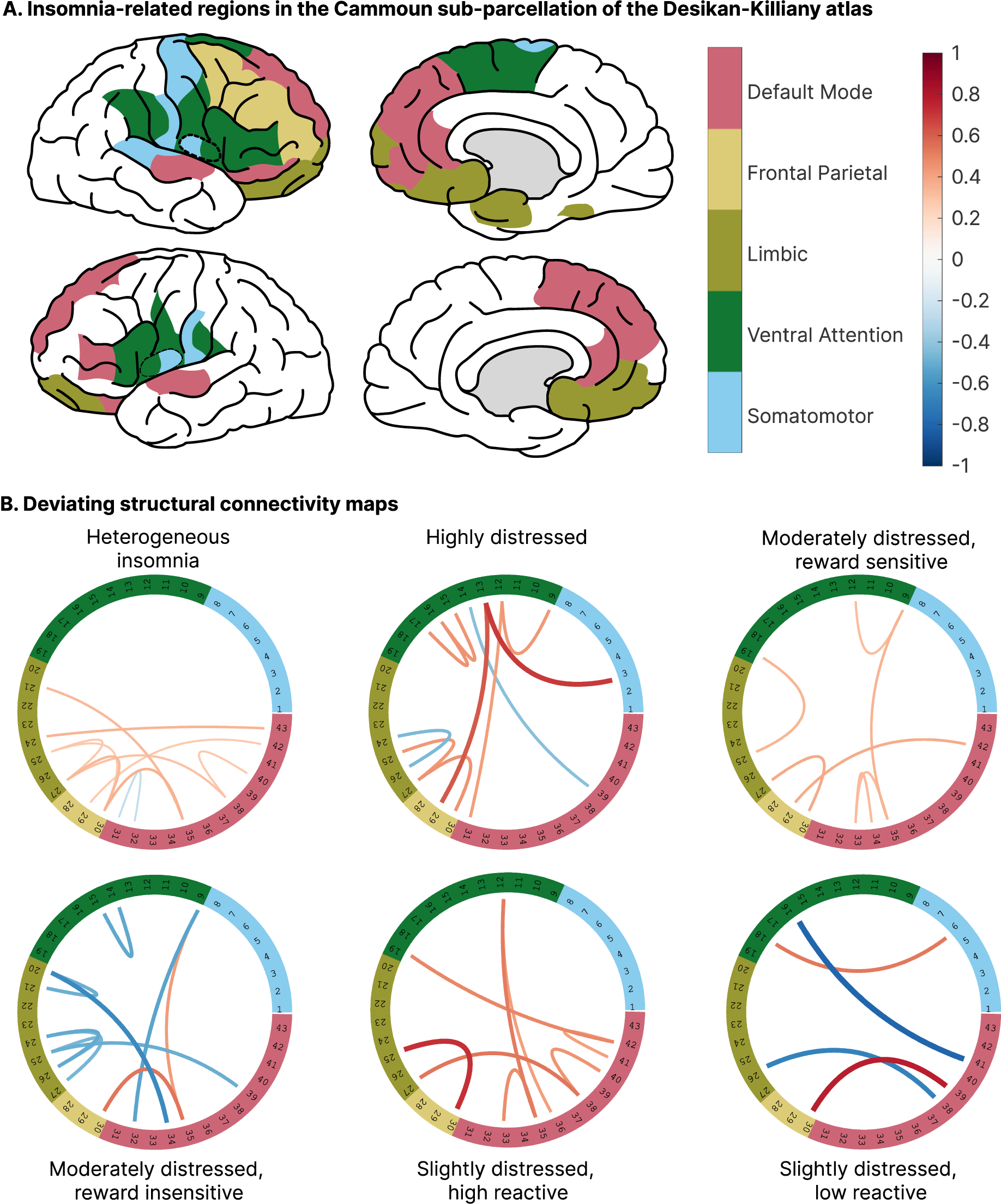
Deviating structural connectivity maps with functional annotation. A) Representation of brain regions of the Cammoun sub-parcellation of the Desikan-Killiany cortical atlas that are, according to the Neuro Knowledge Engine database, involved in insomnia subtype-distinguishing mood and personality traits. Colors indicate the functional networks that the regions belong to according to Yeo *et al.* (45). B) Circular representations of deviating structural connections based on fractional anisotropy. Ring numbers indicate cortical area (table s2) and colors indicate the functional networks (as in A). Thickness and color of the connecting lines represent the standardized effects size of the deviation relative to control, which varied between −0.80 and 0.77.

### Insomnia subtypes differ in structural connectivity deviations

Focusing on the FA-weighted structural connectivity of the selected regions-of-interest relevant to insomnia subtypes, we contrasted people without sleep complaints with heterogeneous insomnia disorder first, and subsequently with each of the five subtypes separately. For both the heterogeneous sample of participants with insomnia as well for the subtypes, deviations in structural connectivity of the 43 regions-of-interest were visualized as red and blue lines, respectively representing higher and lower FA values relative to controls (figure 2B). These deviating connectivity maps indicate differentiating patterns for different subtypes compared to people without sleep complaints, including some opposing deviations. For example, compared to people without sleep complaints, the *moderately distressed reward insensitive* type showed *reduced* connectivity strength between frontal cortical regions, whereas other subtypes showed *increased* connectivity strength.

### Structural connectivity deviations of insomnia subtypes concentrate in different functional networks

To place our findings in a functional perspective we annotated the regions-of-interest in accordance with the functional networks described by Yeo *et al.* (45). The frontal, orbitofrontal and temporal brain regions related to mood and personality traits-distinguishing subtypes were part of five of the seven functional networks described by Yeo *et al.* (45), i.e. the limbic (8 regions), ventral attention (11 regions), somatomotor (8 regions), default mode (13 regions) and frontal parietal network (3 regions, figure 2A). The annotated deviating structural connectivity maps show how involved functional systems differ depending on the subtype of insomnia (figure 2B). To obtain a more globally integrated interpretation of these subtype-specific deviations, we quantified the percentage of brain regions with deviating structural connectivity in each functional network for every group (figure 3). The resulting ‘connectivity deviation profiles’ show that in the heterogeneous total sample, connectivity deviations occur in the limbic, frontal parietal and default mode network. Profiles of the insomnia subtypes revealed differential degrees of involvement of these networks and moreover involvement of the somatomotor network and ventral attention network. Insomnia subtypes differed in how deviations in structural connectivity concentrated in five functional networks. The *highly distressed* subtype distinguished itself from other subtypes in showing more brain regions with structural connectivity deviations in the ventral attention network (7/11 regions, 64%). In the *moderately distressed reward insensitive* subtype connectivity deviations were most concentrated in the limbic network (6/8 regions, 75%). In the *slightly distressed high reactive* subtype, deviations concentrated in the default mode network (7/13 regions, 54%). It should be noted that only 3 of the 43 regions-of-interest for insomnia belong to the frontal parietal system, so that the resolution of perceptual involvement visualized in figure 3 is not optimal for this network. For the absolute number of deviating connections in each functional network we refer to the supplement (see figure s5-8).

**Figure 3.**
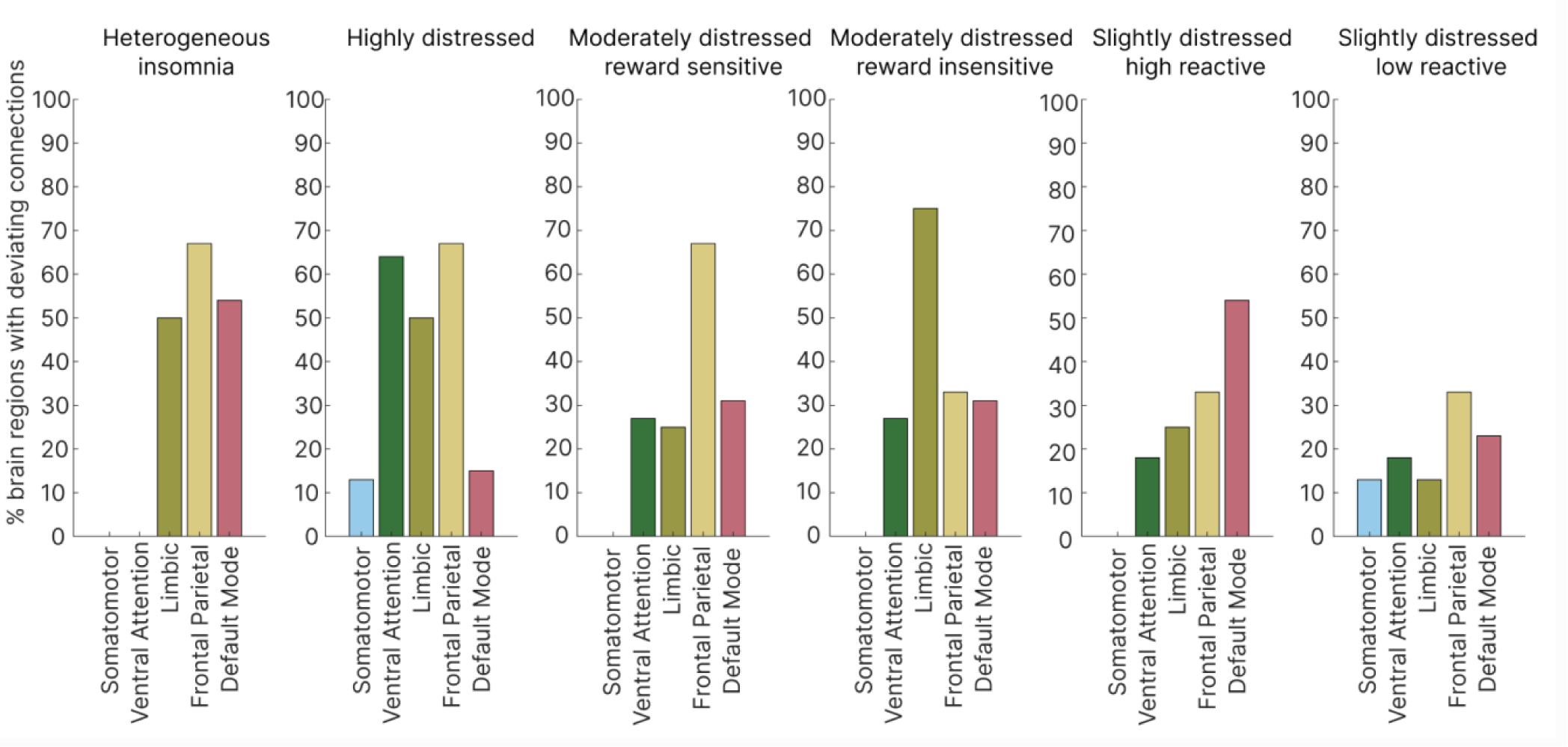
Connectivity deviation profiles. Subtype-specific deviations quantified as the percentage of insomnia-related brain regions with deviating structural connectivity in each functional network. Brain regions defined by the Cammoun sub-parcellation of the Desikan-Killiany cortical atlas. Connectivity deviations concern fractional anisotropy.

### Testing subtype specificity

So far, visual inspection suggests that deviating connectivity is differentially represented in functional systems depending on the subtype of insomnia. To test whether deviating patterns are subtype-specific and would not occur with random sampling from the heterogeneous sample of people with insomnia, we ran 10 000 permutations by shuffling subtype labels across all participants with insomnia while keeping the control labels of the group without sleep complaints the same. We found that the connectivity deviation profile of the *highly distressed* (p = 0.019, p_Bonferroni_ = 0.095) and *moderately distressed reward insensitive* (p = 0.014, p_Bonferroni_ = 0.07) subtypes significantly differed from random subsamples drawn from the heterogeneous pool of mixed insomnia subtypes.

### Subcortical brain regions and validation using a different cortical parcellation

To validate our findings and include subcortical regions, we reconstructed the heat map, repeated the connectivity map analysis and repeated the connectivity deviation profile analysis, now using the 82-parcel Desikan-Killiany combined cortical and subcortical atlas as present in FreeSurfer (42). The heat map analysis selected 38 regions-of-interest resembling the cortical regions selected by the approach using the Cammoun 114-cortical parcel approach, but also subcortical structures, i.e., the left putamen and, bi-lateral hippocampus, amygdala and nucleus accumbens area. The deviating connectivity maps again showed differentiating patterns in different subtypes compared to people without sleep complaints, with sometimes opposing deviations (figure 4B). Visual inspection of the connectivity deviation profiles showed more subtle, but distinct differences between subtypes and indicated involvement of subcortical regions in especially the *moderately distressed reward insensitive* and *slightly distressed low reactive* subtype (figure 5). Permutation testing revealed that profiles of the *highly distressed* (p = 0.018, p_Bonferroni_ = 0.09), *moderately distressed, reward insensitive* (p = 0.002, p_Bonferroni_ = 0.01) and *slightly distressed, low reactive* (p = 0.006, p_Bonferroni_ = 0.03) subtype significantly differed from randomly labeled subsamples drawn from the heterogeneous pool of mixed insomnia subtypes.

**Figure 4.**
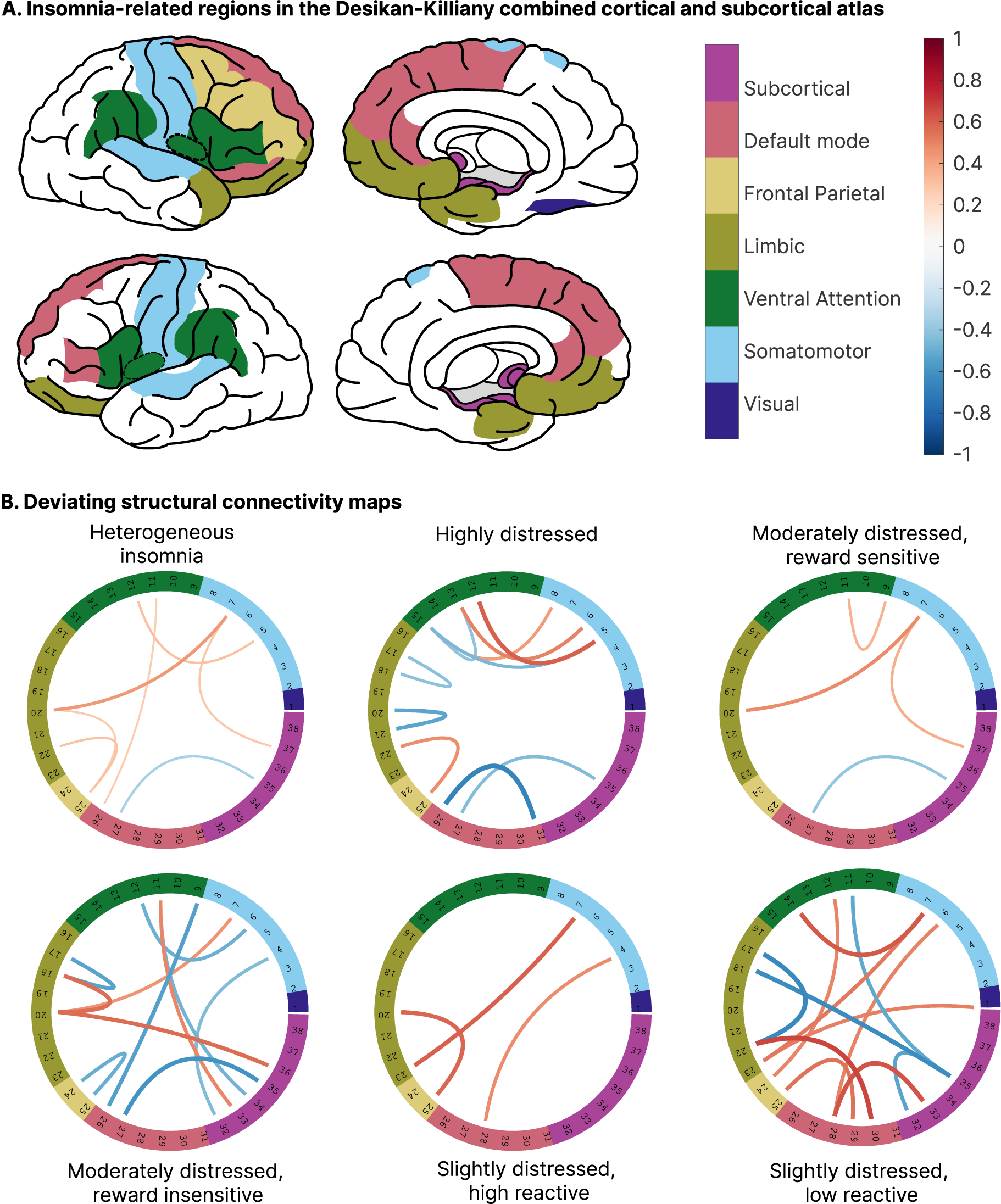
Deviating structural connectivity maps within functional networks including subcortical regions. A) Representation of brain regions of the Desikan-Killiany combined cortical and subcortical atlas that are, according to the Neuro Knowledge Engine database involved in insomnia subtype-distinguishing mood and personality traits. Colors indicate the functional networks that the regions belong to according to Yeo *et al.* (45). B) Circular representations of deviating structural connections based on fractional anisotropy. Ring numbers indicate cortical area (table s3) and colors indicate the functional networks (as in A). Thickness and color of the lines represent the standardized effects size of the deviation relative to control, which varied between −0.67 and 0.67.

**Figure 5.**
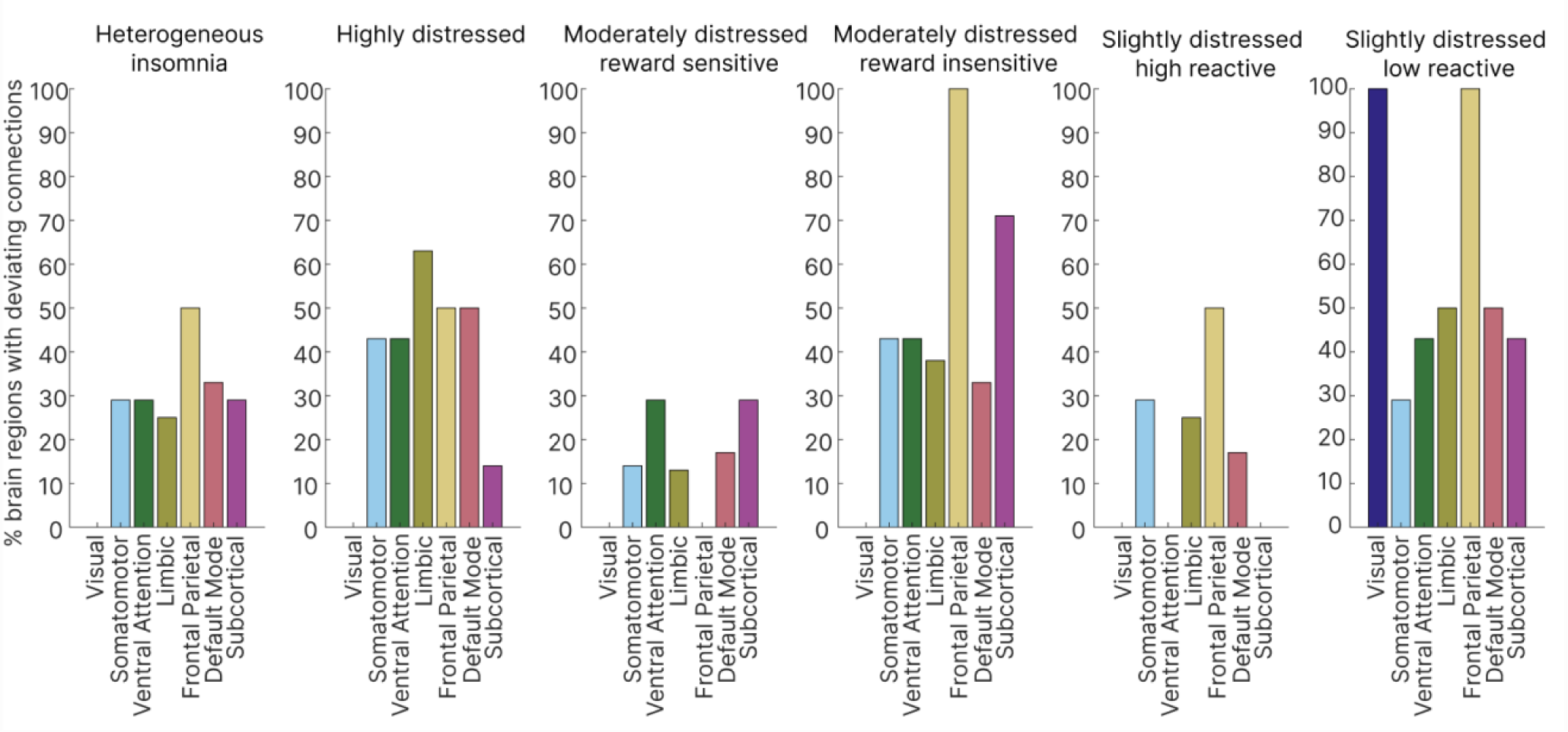
Connectivity deviation profiles including subcortical regions. Subtype-specific deviations quantified as the percentage of insomnia-related brain regions with deviating structural connectivity in each functional network. Brain regions defined by the Desikan-Killiany combined cortical and subcortical atlas. Connectivity deviations concern fractional anisotropy.

### Mean diffusivity-weighted connectivity

In addition to the FA-weighted structural connectivity findings reported on above, we also analyzed MD-weighted structural connectivity. In both atlases, deviating connectivity maps showed more connections with predominantly lower MD in insomnia than in people without sleep complaints. Figure s2 shows that distinctive patterns of deviating connectivity of subtypes compared to people without sleep complaints were once more observed. Permutation testing indicated that the connectivity deviation profile of the *slightly distressed high reactive* subtype significantly differed from random subsamples of heterogeneous insomnia (Cammoun sub-parcellation of the Desikan-Killiany cortical atlas p = 0.006, p_Bonferroni_ = 0.03; Desikan-Killiany combined cortical and subcortical atlas p = 0.020, p_Bonferroni_ = 0.1, figure s3).

## Discussion

This study shows that specific subtypes of insomnia have different patterns of deviating structural connectivity. Deviations in structural connectivity associated with insomnia disorder differ depending on the specific subtype of insomnia, and may even be in opposing directions. For regions-of-interest defined by the circuits associated with subtype-distinguishing personality and mood traits, deviating structural connectivity was predominantly found in limbic, default mode and ventral attention networks. The extent to which brain regions within these networks showed deviating structural connectivity differed between subtypes. We demonstrated robustness of these findings across two brain atlases and by permutation testing. Our findings indicate that insomnia subtypes have distinct profiles of structural connectivity deviations, some of which would dilute or even cancel out in insomnia samples that are heterogeneous with unknown proportional representation of the different subtypes.

### Heterogeneous insomnia disorder

Consistent across two atlases, the heterogeneous insomnia sample predominantly showed deviating connectivity of frontal regions with limbic, default mode and frontal parietal networks. Previous studies in heterogeneous samples of people with insomnia disorder have reported similar deviating structural connectivity in regions such as the orbital frontal gyrus, superior frontal gyrus, cingulate gyrus and, as well as the limbic and default mode network (30,48–50). It is important to stress that the current study focused on finding differences in connectivity deviations between subtypes, and may therefore have missed connectivity deviations that may be generic for all people with insomnia, irrespective of subtype. For example, the angular gyrus was not part of the regions-of-interest we defined based on subtype-distinguishing personality and mood traits, hence we did not evaluate a previously suggested deviation of angular gyrus connectivity (30). Previously reported altered connectivity of the insula (48–50), salience network and ventral attention network (14,16,17) were only observed in the low-resolution combined cortical and subcortical atlas of Desikan-Killiany.

Differences with previous studies may thus partly be related to the regions-of-interest approach we applied, but could also result from different distributions of subtypes within study samples. For example, while diluted and not significant in our larger but heterogeneous total sample, subtype samples showed specific deviating structural connectivity of the insula and other regions linked to the ventral attention network. These findings suggest that the unknown differences in the proportional representation of subtypes in heterogeneous insomnia samples could have contributed to inconsistent findings (19,20).

### Insomnia subtypes

For four out of five insomnia subtypes, their specific connectivity deviation profile based on either FA or MD, differed significantly from profiles of randomly labeled subsamples of heterogeneous insomnia. Subtypes differed with respect to the predominantly affected functional network. In the *highly distressed* subtype, deviating structural connectivity predominantly concentrated in brain regions linked to the ventral attention and limbic network. In the *moderately distressed reward insensitive* subtype deviations predominantly concerned connectivity of subcortical regions and the limbic network. In the *slightly distressed low reactive* subtype deviations predominantly concerned connectivity of the limbic and default mode network. Finally, the *slightly distressed high reactive* subtype was characterized by deviating MD-weighted connectivity in the somatomotor network, ventral attention network, limbic network and default mode network. Our results show, for at least four out of five insomnia subtypes, that connectivity deviation profiles are specific and diluted in randomly labeled subsamples of heterogeneous insomnia. The findings support the notion of different neural correlates underlying insomnia subtypes.

### Functional networks

Functional annotation showed that deviations in structural connectivity concentrated in the ventral attention network, limbic network, default mode network and subcortical regions which have all previously been linked to insomnia disorder or traits relevant to insomnia. The **ventral attention network** described by Yeo *et al.* is likely an aggregate of the salience and cingulo-opercular networks (45). This network is important in identifying and responding to salient stimuli, as well as shifting attention following unexpected stimuli (51,52). Previous studies have reported deviating structural (30,48) and functional (53) connectivity of the ventral attention network or salience network in insomnia. Altered salience network connectivity has also been reported in other psychiatric disorders (54,55) characterized by insomnia complaints and polygenic risks that overlap strongly with the polygenic risk of insomnia (56,57), notably depression and anxiety. It is tempting to suggest that the marked deviating structural connectivity in the ventral attention network in the *highly distressed* subtype could be associated with their most distinctive high level of pre-sleep arousal (22).

Within the regions of interest defined by their involvement in the personality and mood traits that distinguish insomnia subtypes, the **limbic network** comprised of orbital frontal regions and the right frontal pole. Orbitofrontal brain regions are important in emotion, processing reward value, decision making, and problem-solving abilities (58,59). Alterations in structure and/or function of the orbitofrontal cortex have been described in insomnia (60–64) and depression (59). Several studies even suggest the orbitofrontal cortex as a region where deviations link insomnia and depression (65–67). Interestingly, the *highly distressed* and *moderately distressed reward insensitive* subtype showed a high degree of deviating limbic structural connectivity and is indeed characterized by reduced subjective happiness, reduced positive affect and higher prevalence of depression compared to other subtypes (22). On the other hand, the *slightly distressed low reactive* subtype also showed disturbed limbic connectivity in the Desikan-Killiany combined cortical and subcortical atlas, but is not characterized by similarly excessive deviations in traits characteristic of depression (22). A possible explanation could be that disturbed limbic connectivity is likely to occur in all people suffering from insomnia, while the deviating connections and direction of deviations may differ between subtypes (see Figure 2B and 4B).

Deviating connectivity of brain areas belonging to the **default mode network** was observed in all subtypes to a certain extent, and did not seem to differentiate between subtypes. With its involvement in internal thought (68,69) and maladaptive rumination (70,71), the default mode network is of interest due to its potential role in dysfunctional forms of cognitive control in insomnia (53,72–74). Early life experiences could play an role as they have long-lasting effects on functional connectivity including the default mode network (75,76) and contribute to the risk of developing insomnia (77,78) as well. The observed deviating **subcortical** connectivity is in line with previous described altered frontal-subcortical connectivity (48,49,79). The number of subcortical regions with deviating structural connectivity differed between the five subtypes and was highest in *moderately distressed reward insensitive* subtype.

### Limitations

Limitations have to be considered. First, group sizes were conventional for neuroimaging studies, but the required power for brain-wide association studies remains an ongoing debate (80–83). To avoid underpowered comparisons, we integrated deviating structural connections into connectivity deviation profiles before performing statistical testing. Hence, findings should be interpreted at the profile level. Second, our study focused on a data-driven selection of cortical and subcortical regions. As a result, some regions and connections reported on in previous studies of insomnia were outside our scope. Lastly, we used mean FA and MD of reconstructed tractography streamlines as connectivity strength, while some previous studies used streamline density (30,48,49). Both methods have their limitations (84), but FA and MD measures have proven to be a sensitive proxy to detect changes in white matter microstructure (29).

### Conclusion

Our study provides a first indication that insomnia subtypes show differentiating, subtype-specific structural connectivity deviation profiles. For four out of five subtypes, their subtype-specific structural connectivity deviation profile differed significantly from what can be expected in randomly drawn subsamples of heterogeneous insomnia. Our findings support the notion of insomnia subtypes with possibly different underlying brain mechanisms and shows that subtyping of insomnia could be essential to discover more robust structural brain correlates and gain understanding of brain mechanisms involved. A freely available tool to do so online is currently being developed (https://staging.insomniatype.org/).

## Acknowledgements

This project benefitted from funding by ZonMw, Open Competition project 09120011910032 REMOVE; and from the European Research Council (ERC), Advanced Grant 101055383 OVERNIGHT. TB, JL and OL have been supported by Vrije Universiteit Amsterdam University Research Fellowships.

