## supplement for "Insomnia subtypes have differentiating deviations in brain structural connectivity"

### Supplementary material;

#### Methods

*Exclusion criteria*

Use of sleep medication within the 2 months prior to data acquisition was an exclusion criterium in sample 1,2 and 4 ($N_{insomnia}$ = 44, $N_{control}$ = 38). Sleep medication was allowed and monitored in sample 3 ($N_{insomnia}$=160, $N_{control}$ =35, table s1). In addition, current treatment with antidepressant medication or current cognitive behavioral therapy for insomnia were study specific exclusion criteria in sample 3. Groups did not show significant differences in medication use.

*MRI acquisition*

Participants were scanned on a Philips Achieva 3T at two different locations of the Spinoza Centre for Neuroimaging in Amsterdam, The Netherlands (location Meibergdreef: sample 1 $N_{insomnia}$=7, $N_{control}$ =4; sample 2 $N_{insomnia}$=13, $N_{control}$ =8; sample 3 $N_{insomnia}$=160, $N_{control}$ =35; location Roeterseiland: sample 4; $N_{insomnia}$=24, $N_{control}$ =26). Participants were asked to refrain from alcohol and drugs on assessment days and were not allowed to consume caffeinated beverages at least six hours before MRI assessments. Head motion during scanning was restricted by foam pads. T1-weighted images for sample 1, 2, and 4 were acquired with a 3D Turbo Field Echo sequence and the following scanning parameters: repetition time (TR)/ echo time (TE) = 8.3/3.8 ms, voxel size = $(1mm)^3$, 220 slices, field of view (AP × RL × FH) = 240 × 188 × 220mm, flip angle = 8°, encoding direction = RL. For sample 3, T1-weighted images were acquired according to the ADNI protocol (1): TR/ TE = 6.5/2.9ms, voxel size = $(1mm)^3$, 211 slices, field of view (AP × RL × FH) = 256x256x211mm, flip angle = 9°, encoding direction = anterior-posterior. All diffusion weighted images were acquired using a 32-channel head coil. For sample 1, 2, and 4 a single-shot echo-planar imaging sequence was used with 32 diffusion gradient directions (b=1000 sec/mm2) followed by one additional image without diffusion weighting (b=0 sec/mm2). Scanning parameters were: TR/ TE = 16070/48ms (shortest), voxel size = $(2mm)^3$, 65 slices without gap covering the whole brain, field of view

(AP × RL × FH) = 224 x 224 x 130mm, flip angle = 90°, encoding direction = anterior-posterior (right-left for sample 1). For sample 3 diffusion weighted images were acquired using a multiband spin-echo echo-planar imaging (EPI) sequence with 9 non-diffusion weighted (b=0) images and 88 diffusion weighted images (29 with b=1000 s/mm$^2$, 59 with b=2000 s/mm$^2$). Scanning parameters were: TR/ TE = 4683/95ms (shortest), voxel size = (2mm)$^3$, 66 slices without gap covering the whole brain, field of view (AP × RL × FH) = 224 x 224 x 132mm, flip angle = 90°, encoding direction = anterior-posterior, multiband SENSE = 2. In addition, we acquired one non-diffusion weighted scan in the opposite phase encoding direction (posterior-anterior) to estimate susceptibility-induced distortions.

*Structural connectome reconstruction*

Preprocessing and reconstruction of the structural connectivity networks was performed separately for each sample. T1w scans were preprocessed and segmented using 'recon-all' function of FreeSurfer (2) stable version 6.0.1. The segmentation and original T1w scan were used to preprocess the diffusion weighted data with the Connectivity Analysis TOolbox (3) (CATO, version 3.1.2) and the FMRIB Software Library (4) (FSL version 6.0.4). CATO configuration included the correction of the diffusion weighted data for eddy currents, subject movement and susceptibility induced distortions using 'eddy' (5). In sample 3 a non-diffusion weighted scan with opposite phase encoding direction was available to correct for susceptibility induced distortions by combining 'eddy' with 'topup'(5,6). Subsequently CATO reconstructed the diffusion using the Diffusion Tensor Imaging (DTI) model combined with Constrained Spherical Deconvolution (CSD)and performed fiber tracking.. For the network reconstruction we used the fine-grained Cammoun sub-parcellation of the Desikan-Killiany atlas consisting of 114 cortical regions (3,7). To validate our findings and include subcortical regions, we also used the Desikan-Killiany combined cortical and subcortical atlas as present in FreeSurfer (8) consisting of 82 regions. For each atlas, the weight of a connection between regions was computed as the weighted average FA or MD value over all voxels that streamlines passed. Quality control was applied by filtering the connectivity matrices for outliers. Extreme outliers were defined as matrices of which the mean number of streamlines, mean FA, mean MD, average prevalence of present connections, average prevalence of absent connections and the number of connected regions of a subject were 3 times the interquartile range above the $3^{rd}$ quartile or below the $1^{st}$ quartile and excluded from analysis. To ensure sufficient data points for each connection, analyses included only connections that were present in at least 70% of the subjects (9).

*Data harmonization*

To account for multisite and multi-sample effects we harmonized the connection weights between all brain regions for sample differences using the 'NeuroCombat' R-package version 1.0.13 (https://github.com/Jfortin1/ComBatHarmonization) in R(10) version 4.0.4. To preserve biological variance of interest we included a covariate matrix including group (control and insomnia subtype labels), age, sex and brain volume. Sample 3 was set as reference dataset given the large number of participants in that sample and the additional correction for distortions using FSL 'topup'. Because unmeaningful zeros influence the harmonization procedure, we converted all connections with a zero value to a missing value (NaN, not-a-number) before harmonizing the connectivity data. Subsequent analyses used the harmonized connectivity values.

### Results

#### Structural connectivity weighted by FA

Examining the deviations exceeding a primary (t-statistic) threshold of |t| >= 2 in the Cammoun sub-parcellation of the Desikan-Killiany cortical atlas, heterogeneous insomnia disorder (n=204) differed from good sleepers (n=73) in 9 connections with slightly higher FA (β = 0.275 to 0.356) and 1 connection with lower FA (β = -0.260). These deviating connections mainly occurred within and between regions of the default mode network and limbic network.

Compared to good sleepers, the *highly distressed subtype* (n=31) had 8 connections with higher FA (β =0.424 to 0.694) and 2 connections with lower FA (β = -0.452, -0.423). These were predominant within and between two regions of the frontal parietal network, ventral attention network and limbic network.

The *moderately distressed, reward sensitive subtype* (n=89) had 7 connections with higher FA (β =0.321 to 0.383). These sparsely occurred between the default mode network, frontal parietal network, limbic network and ventral attention network.

The *moderately distressed, reward insensitive subtype* (n=30) had 2 connections with higher FA (d= 0.431, 0.538) and 8 connections with lower FA (β =-0.638 to -0.464). These deviations were concentrated within the limbic network and in connections between several resting state networks with the default mode network.

The *slightly distressed, high reactive subtype* (n=34) had 8 connections with higher FA (β = 0.412 to 0.719). These deviations were predominantly within the default mode network and in connections with the limbic network and ventral attention network.

The *slightly distressed, low reactive subtype* (n=20) had 2 connections with higher FA (d= 0.530, 0.771) and 2 connections with lower FA (β =-0.797, -0.675). These deviations occurred mainly between the default mode network and frontal parietal network, limbic network, and ventral attention network.

#### Structural connectivity weighted by MD

Examining the deviations exceeding a primary (t-statistic) threshold of |t| >= 2 in the Cammoun sub-parcellation of the Desikan-Killiany cortical atlas, heterogeneous insomnia disorder differed from good sleepers in 32 connections with moderately lower MD (β = -0.455 to -0.264). These deviating connections occurred across the default mode network, frontal parietal network, limbic network, somatomotor network and ventral attention network.

Compared to good sleepers, the *highly distressed subtype* had 20 connections with lower MD (β = -0.719 to -0.406). These deviating connection occurred across the default mode network, frontal parietal network, limbic network, somatomotor network and ventral attention network.

The *moderately distressed, reward sensitive subtype* had 17 connections with lower MD (β = -0.530 to -0.304). These predominantly occurred within and between the default mode network and limbic network.

The *moderately distressed, reward insensitive subtype* (n=30) had 1 connection with higher MD (β = 0.699) and 6 connections with lower MD (β =-0.626 to -0.435). These sparse deviations were occurred between the default mode network, somatomotor network and ventral attention network.

The *slightly distressed, high reactive subtype* had 1 connection with higher MD (β = 0.515) and 43 connections with lower MD (β =-0.733 to -0.408). These deviating connections occurred across the default mode network, frontal parietal network, limbic network, somatomotor network and ventral attention network.

The *slightly distressed, low reactive subtype* had 12 connections with lower MD (β =-0.833 to -0.508). These deviations occurred mainly within limbic network and ventral attention network.

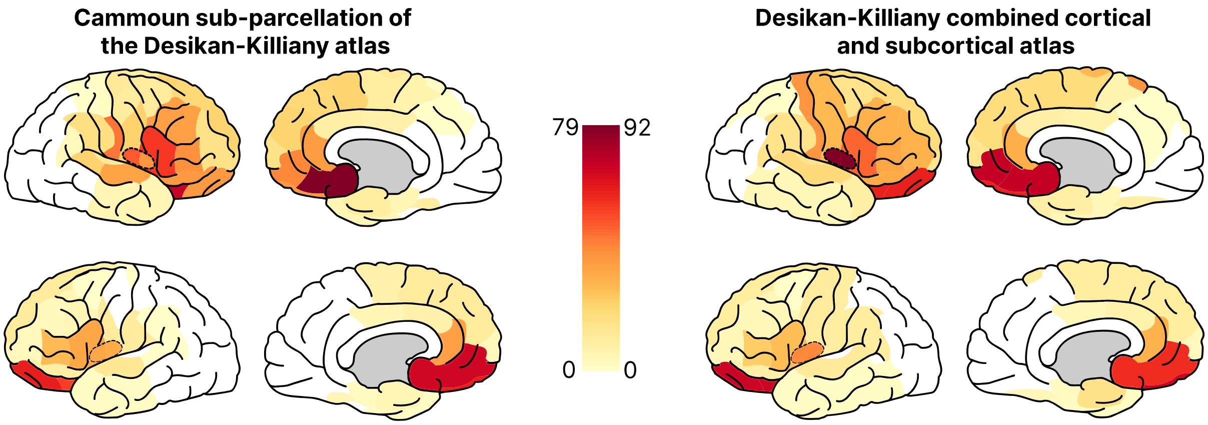


**Figure s1. Heat map of regions implicated in insomnia subtype-defining traits.**

Heat maps of the extent to which brain areas have been implicated in subtype-distinguishing personality and mood traits as represented by the terms linked to insomnia subtype-distinguishing personality and mood traits and listed below.

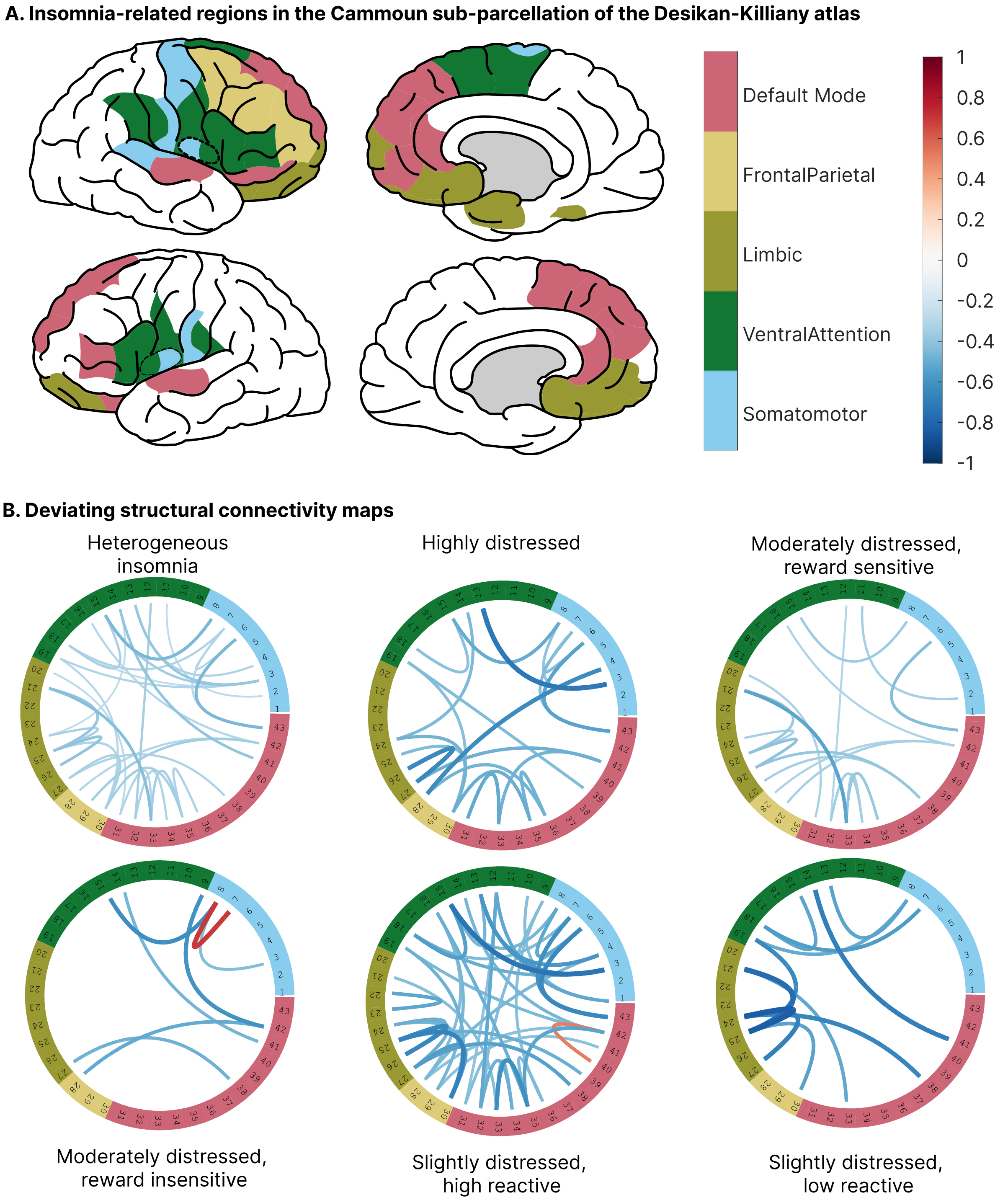


**Figure s2. Deviating structural connectivity maps within functional networks based on MD.** A) Representation of brain regions of the Cammoun sub-parcellation of the Desikan-Killiany cortical atlas that are, according to the Neuro Knowledge Engine database involved in insomnia subtype-distinguishing mood and personality traits. Colors indicate the functional networks that the regions belong to according to Yeo *et al.* (11). B) Circular representations of deviating structural connections based on mean diffusivity. Ring numbers indicate cortical area (table s2) and colors indicate the functional networks (as in A). Thickness and color of the lines represents the standardized effects size of the deviation relative to control, which varied between -0.26 and -0.83.

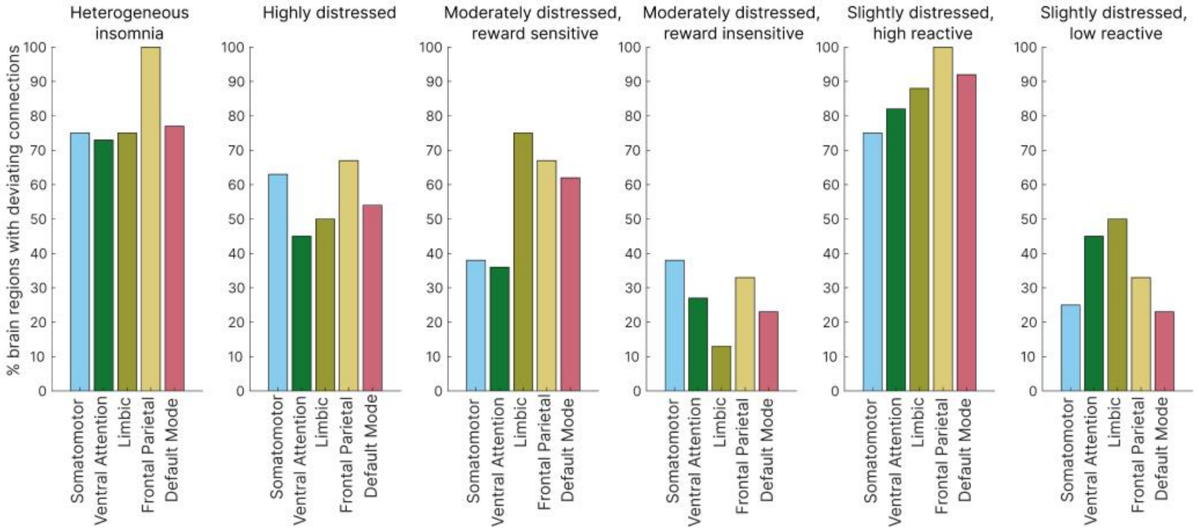


**Figure s3. Connectivity deviation profiles based on MD.**

Subtype-specific deviations quantified as the percentage of insomnia-related brain regions with deviating structural connectivity in each functional network. Brain regions defined by the Cammoun sub-parcellation of the Desikan-Killiany cortical atlas. Connectivity deviations concern mean diffusivity.

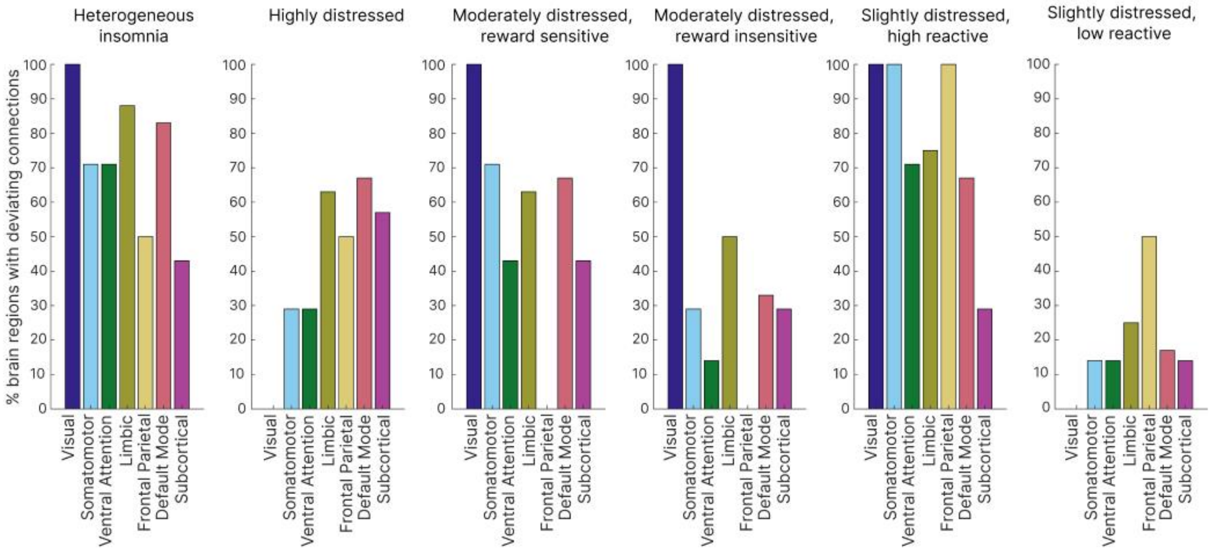


**Figure s4. Connectivity deviation profiles including subcortical regions based on MD.** Subtype-specific deviations quantified as the percentage of insomnia-related brain regions with deviating structural connectivity in each functional network. Brain regions defined by the Desikan-Killiany combined cortical and subcortical atlas. Connectivity deviations concern mean diffusivity.

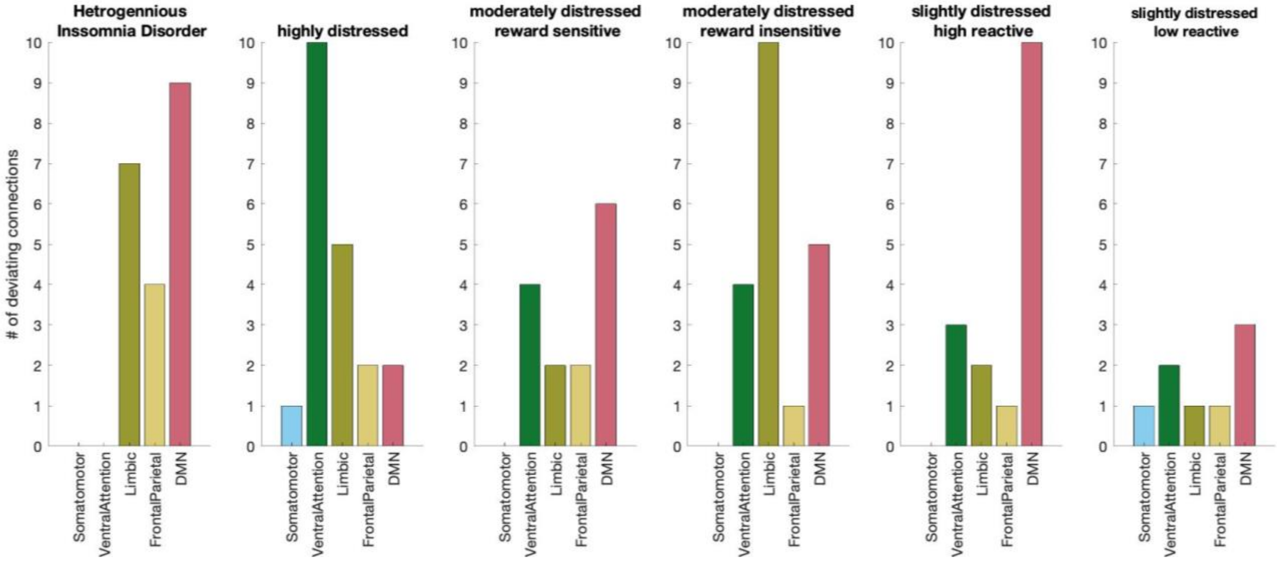


**Figure s5. Absolute connectivity deviation profiles based on FA.**

Subtype-specific deviations quantified as the total number of deviating connections based on fractional anisotropy (FA) in each functional network. Brain regions defined by the Cammoun sub-parcellation of the Desikan-Killiany atlas.

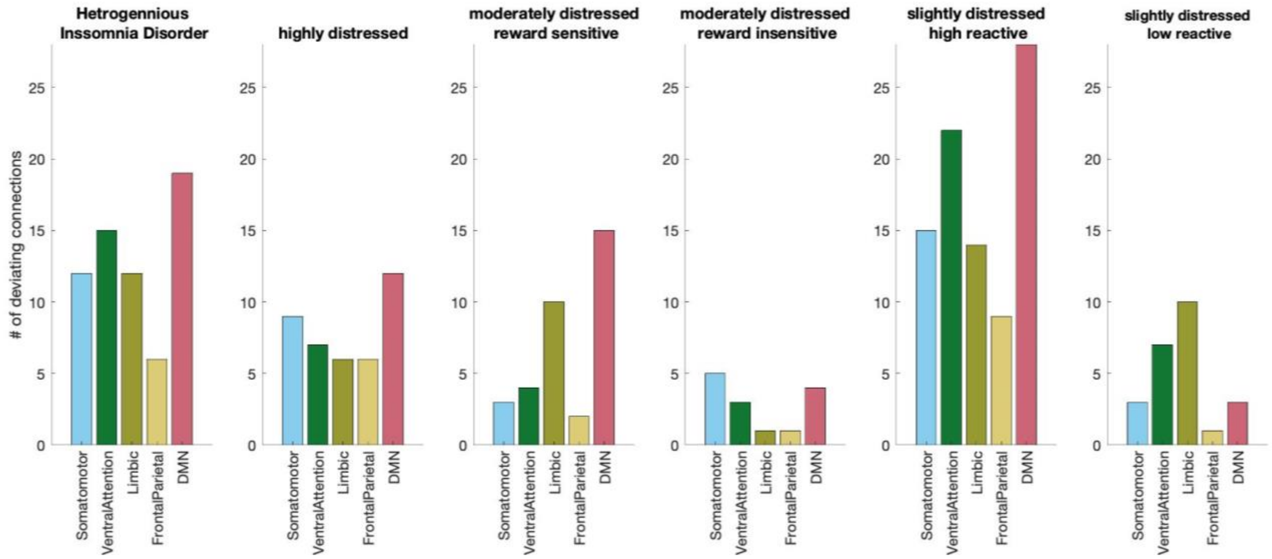


**Figure s6. Absolute connectivity deviation profiles based on MD.**

Subtype-specific deviations quantified as the total number of deviating connections based on mean diffusivity (MD) in each functional network. Brain regions defined by the Cammoun sub-parcellation of the Desikan-Killiany atlas.

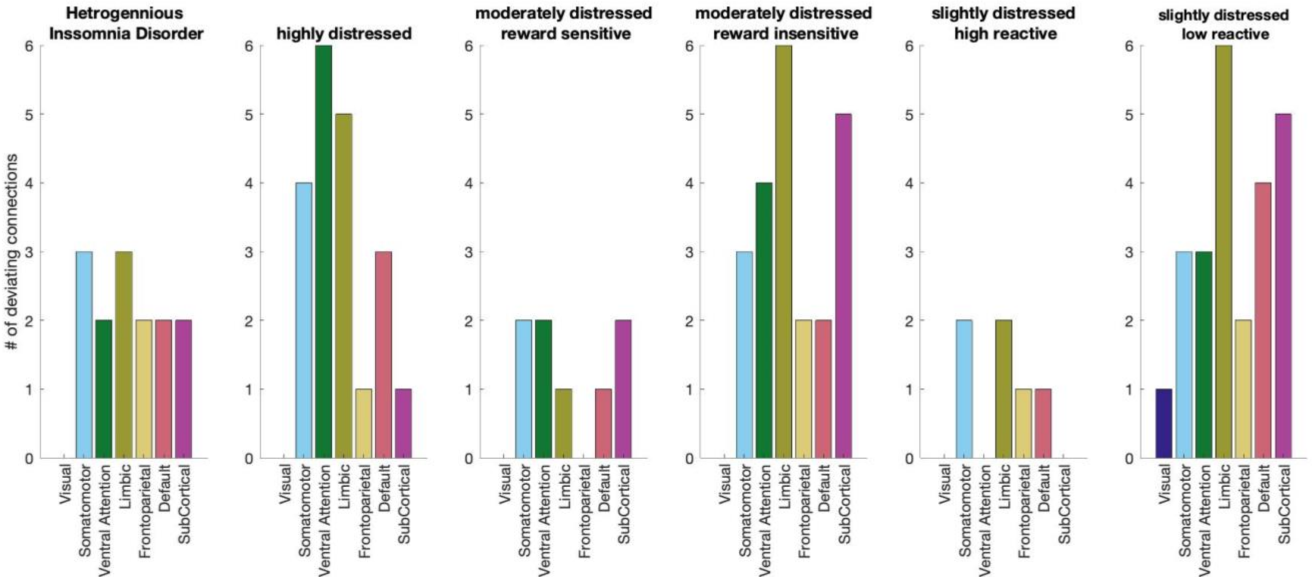


**Figure s7. Absolute connectivity deviation profiles including subcortical regions based on FA.** Subtype-specific deviations quantified as the total number of deviating connections based on fractional anisotropy (FA) in each functional network. Brain regions defined by the Desikan-Killiany combined cortical and subcortical atlas.

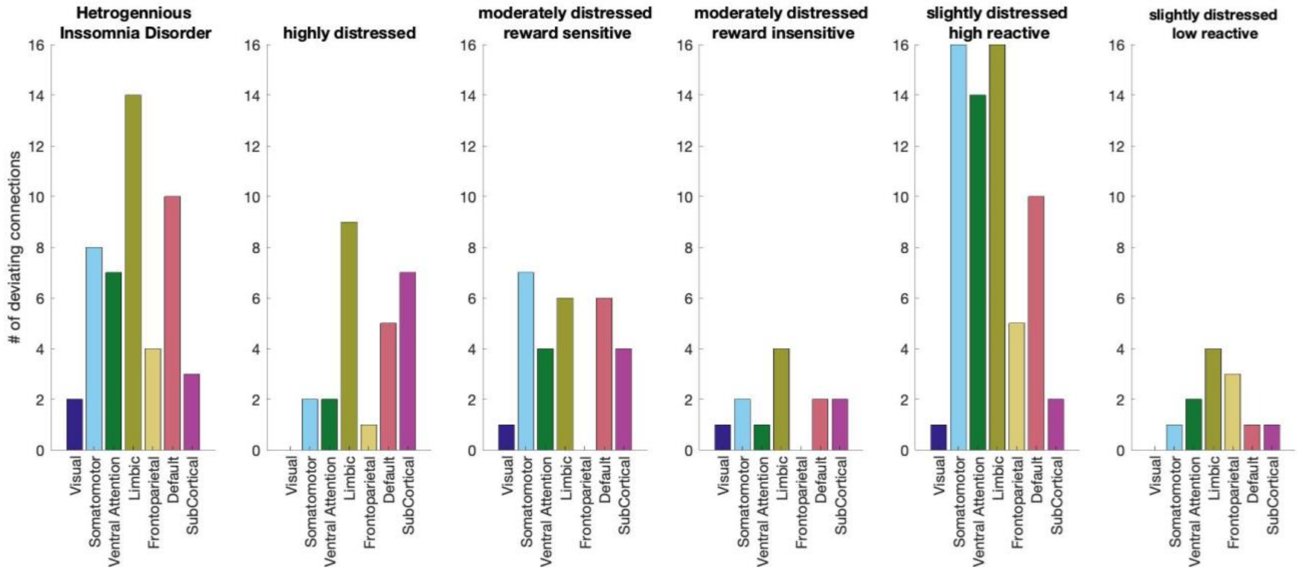


**Figure s8. Absolute connectivity deviation profiles including subcortical regions based on MD.** Subtype-specific deviations quantified as the total number of deviating connections based on mean diffusivity (MD) in each functional network. Brain regions defined by the Desikan-Killiany combined cortical and subcortical atlas.

**Table s1.** Overview of medication use in sample 3.

| | | Insomnia disorder (N=160) | | | | | |
|---|---|---|---|---|---|---|---|
| | **Control** | **Highly distressed** | **Moderately distressed reward sensitive** | **Moderately distressed reward insensitive** | **Slightly distressed high reactive** | **Slightly distressed low reactive** | p |
| n | 35 [a] | 26 | 72 | 28 | 20 | 14 | |
| Sleep medicines | 0 | 3 | 12 | 2 | 1 | 1 | 0.131 |
| Antidepressants | 1 | 0 | 1 | 0 | 0 | 0 | 0.817 |
| Anti-anxiety drugs | 0 | 0 | 1 | 0 | 0 | 0 | 0.888 |
| Anti-psychotics | 0 | 0 | 0 | 0 | 0 | 0 | - |
| Antihypertensives | 1 | 1 | 8 | 2 | 1 | 2 | 0.569 |
| Thyroid medication | 1 | 0 | 2 | 1 | 1 | 1 | 0.865 |
| Antiasthmatics | 1 | 3 | 5 | 3 | 2 | 0 | 0.624 |
| Anti-Parkinson drugs | 0 | 0 | 0 | 0 | 0 | 0 | - |
| Anticonvulsants | 0 | 0 | 0 | 0 | 1 | 0 | 0.12 |
| Headache medicines | 2 | 0 | 4 | 2 | 1 | 1 | 0.874 |
| Stimulants | 0 | 1 | 1 | 0 | 0 | 0 | 0.683 |
| Others | 6 | 7 | 12 | 7 | 6 | 3 | 0.726 |

[a] Data on medication use was unavailable for n = 1.

**Table s2.** Mapping of the Cammoun sub-parcellation of the Desikan-Killiany cortical atlas labels to the numbers in the circular representations of the deviating connections in figure 2 and s2.

| Nr. | Cortical region | Nr. | Cortical region |
|---|---|---|---|
| 1 | ctx-lh-postcentral_3 | 23 | ctx-rh-fusiform_2 |
| 2 | ctx-lh-insula_1 | 24 | ctx-rh-lateralorbitofrontal_1 |
| 3 | ctx-rh-postcentral_1 | 25 | ctx-rh-lateralorbitofrontal_2 |
| 4 | ctx-rh-precentral_2 | 26 | ctx-rh-medialorbitofrontal_2 |
| 5 | ctx-rh-precentral_3 | 27 | ctx-rh-frontalpole_1 |
| 6 | ctx-rh-superiortemporal_1 | 28 | ctx-rh-caudalmiddlefrontal_1 |
| 7 | ctx-rh-transversetemporal_1 | 29 | ctx-rh-rostralmiddlefrontal_1 |
| 8 | ctx-rh-insula_1 | 30 | ctx-rh-rostralmiddlefrontal_2 |
| 9 | ctx-lh-parsopercularis_1 | 31 | ctx-lh-lateralorbitofrontal_1 |
| 10 | ctx-lh-precentral_4 | 32 | ctx-lh-parstriangularis_1 |
| 11 | ctx-lh-supramarginal_1 | 33 | ctx-lh-rostralanteriorcingulate_1 |
| 12 | ctx-lh-insula_2 | 34 | ctx-lh-superiorfrontal_1 |
| 13 | ctx-rh-parsopercularis_1 | 35 | ctx-lh-superiorfrontal_2 |
| 14 | ctx-rh-parstriangularis_1 | 36 | ctx-lh-superiorfrontal_3 |
| 15 | ctx-rh-precentral_1 | 37 | ctx-lh-superiortemporal_2 |
| 16 | ctx-rh-superiorfrontal_3 | 38 | ctx-rh-medialorbitofrontal_1 |
| 17 | ctx-rh-superiorfrontal_4 | 39 | ctx-rh-parsorbitalis_1 |
| 18 | ctx-rh-supramarginal_2 | 40 | ctx-rh-rostralanteriorcingulate_1 |
| 19 | ctx-rh-insula_2 | 41 | ctx-rh-superiorfrontal_1 |
| 20 | ctx-lh-lateralorbitofrontal_2 | 42 | ctx-rh-superiorfrontal_2 |
| 21 | ctx-lh-medialorbitofrontal_1 | 43 | ctx-rh-superiortemporal_2 |
| 22 | ctx-rh-entorhinal_1 | | |

**Table s3.** Mapping of the Desikan-Killiany combined cortical and subcortical atlas labels to the numbers in the circular representations of the deviating connections in figure 4.

| Nr. | (Sub)cortical region | Nr. | (Sub)cortical region |
|---|---|---|---|
| 1 | ctx-rh-fusiform | 20 | ctx-rh-lateralorbitofrontal |
| 2 | ctx-lh-postcentral | 21 | ctx-rh-medialorbitofrontal |
| 3 | ctx-lh-precentral | 22 | ctx-rh-frontalpole |
| 4 | ctx-lh-superiortemporal | 23 | ctx-rh-temporalpole |
| 5 | ctx-rh-postcentral | 24 | ctx-rh-caudalmiddlefrontal |
| 6 | ctx-rh-precentral | 25 | ctx-rh-rostralmiddlefrontal |
| 7 | ctx-rh-superiortemporal | 26 | ctx-lh-parstriangularis |
| 8 | ctx-rh-transversetemporal | 27 | ctx-lh-rostralanteriorcingulate |
| 9 | ctx-lh-parsopercularis | 28 | ctx-lh-superiorfrontal |
| 10 | ctx-lh-supramarginal | 29 | ctx-rh-parsorbitalis |
| 11 | ctx-lh-insula | 30 | ctx-rh-rostralanteriorcingulate |
| 12 | ctx-rh-parsopercularis | 31 | ctx-rh-superiorfrontal |
| 13 | ctx-rh-parstriangularis | 32 | Left-Putamen |
| 14 | ctx-rh-supramarginal | 33 | Left-Hippocampus |
| 15 | ctx-rh-insula | 34 | Left-Amygdala |
| 16 | ctx-lh-entorhinal | 35 | Left-Accumbens-area |
| 17 | ctx-lh-lateralorbitofrontal | 36 | Right-Hippocampus |
| 18 | ctx-lh-medialorbitofrontal | 37 | Right-Amygdala |
| 19 | ctx-rh-entorhinal | 38 | Right-Accumbens-area |

**Terms linked to insomnia subtype-distinguishing personality and mood traits**

Listed are the terms related to insomnia-relevant subtype-distinguishing mood and personality traits as used in Blanken *et al.* (12). These terms were matched to keywords present in the Neuro Knowledge Engine database to determine brain regions involved in insomnia subtype-distinguishing mood and personality traits (see Methods).

action_control
affect
agreeableness
arousal
behavioral_activation
childhood_trauma
chronotype
cognition
cognitive
conscientiousness
dampening_of_postivie_moods
duraton_of_insomnia_response_to_life_events
emotion
emotional_memory
experiencing_pleasure
extraversion
fatigue
gonogo_task
happiness
heat/activity_induced_fatigue
insomnia
insomnia_age_of_onset
insomnia_in_family
insomnia_response_to_stress
interoception
life_events
loss
mood
negative_affect
negative_emotion
neuroticism
openness
perfectionism
pleasure
positive_affect
positive_rumination
pre-sleep_arousal
response_inhibition
reward
rumination
sadness
salience
severity_of_insomnia_response_to_life_events
sleepiness
stress
subjective_happiness
trauma
